# Characterization of functional diversity of the human brain based on intrinsic connectivity networks

**DOI:** 10.1101/475855

**Authors:** Congying Chu, Lingzhong Fan, Tianzi Jiang

## Abstract

Spontaneous fluctuations underlying the brain activity can reflect the intrinsic organization of the system, such as the functional brain networks. In large scale, a network perspective has emerged as a new avenue to explore the functional properties of human brain. Here, we studied functional diversity in healthy subjects based on the network perspective. We hypothesized that the patterns of participation of different functional networks were related with the functional diversity of particular brain regions. Independent component analysis (ICA) was adopted to detect the intrinsic connectivity networks (ICNs) based on the data of resting-state functional MRI. An index of functional diversity (FD index) was proposed to quantitatively describe the degree of anisotropic distribution related with participation of various ICNs. We found that FD index continuously varied across the brain, for example, the primary motor cortex with low FD value and the precuneus with significantly high FD value. The FD values indicated the different functional roles of the corresponding brain regions, which were reflected by the various patterns of participation of ICNs. The FD index can be used as a new approach to quantitatively characterize the functional diversity of human brain, even for the changed functional properties caused by the psychiatric disorders.

## Introduction

The functional organization of human brain involves the coordination of brain regions that exhibit various functional properties (Bressler 1995; Bressler and Menon 2010; Felleman and Van Essen 1991; Goldmanrakic 1988; Pessoa 2014; Power et al. 2011; Sporns 2013, 2014). Functional diversity of individual brain area is an important kind of property to systematically understand the large-scale functional organization of human brain. As we know, some brain regions, such as the primary sensory-motor cortices, are observed to participate in specialized functions, such as basic sensory and motor functions (Kanwisher 2010). In particular, visual system has been deep studied to find several separate regions exhibiting highly specialized function such as motion, color and object processing (Anderson et al. 2013; Kanwisher 2010; Zeki 1993). Meanwhile, there are also brain regions required by performing multiple cognitive functions. For example, supramarginal gyrus and middle temporal gyrus are found to be activated across different types of tasks involved with attention and language production (Braga et al. 2013). The various level of functional diversity manifested in human brain reveals the different functional role of brain region. It has become an increasing concern about how to characterize the functional diversity involved with different brain regions.

Thanks to the advances in neuroimaging techniques, especially the functional MRI (fMRI), our understanding about functional diversity of human brain has been advanced. Based on the fast accumulation of task-based fMRI studies, there are some available database with well-collected details, such as BrainMap and Neurosynth (Fox et al. 2005; Fox and Lancaster 2002; Laird et al. 2005; Yarkoni et al. 2011). Through mining these databases, the reverse inferences can be drawn to deduce the engagement of particular functions given the activation in special brain regions (Poldrack 2006, 2011). Further, the functional diversity or flexibility across cerebral cortex was estimated based on BrainMap database (Anderson et al. 2013; Yeo et al. 2014). The main idea behind these two studies is to map the voxel-wise pattern of participating in various task domains or cognitive components. However, the functional diversity estimated from these databases should be considered as a summary based on various task-based studies. As the experimental conditions, including the number of subjects, the experimental paradigm and the detail about collecting / analyzing data, are not exactly the same, it is hard to generalize the results to a particular group of subjects or even to a subject. Based on resting-state fMRI, the overlap of functional networks has been used as an approach to describe the potential functional heterogeneity of cortical regions (Xu et al. 2015; Yeo et al. 2013). However, the number of overlapped functional networks in brain regions is correlated with the statistical threshold as described by Xu et al. (2015). Obviously, a strict threshold will lead to little overlap between functional networks. In addition, the functional diversity reflecting the underlying organization of human brain should be a considered as a continuous property across the whole brain. It might be inappropriate to describe the functional diversity using discrete number, such as the number of functional networks. Further, if we intend to find the change of functional diversity, supposing the existence of the change, between the healthy subjects and the subjects with psychiatric disorder, a quantitative characterization of functional diversity is required. Nevertheless, it is still an open question about how to quantitatively characterize the functional diversity across the whole brain.

In the present study, we sought to provide a frame to characterize the functional diversity via a quantitative approach. As described by Bressler and Menon (2010), the human brain is intrinsically organized as functional networks. More remarkable, ICNs detected in resting-state are generally considered to reflect the networks of brain function, even related with the task-based co-activation networks (Sadaghiani and Kleinschmidt 2013; Smith et al. 2009; Spreng et al. 2010). Using the functional networks as the object of study has opened a new avenue to studies the functional diversity across the cerebral cortex. Based on the viewpoint of network, we hypothesized that a brain region would present a high level of functional diversity if the region significantly involved in various ICNs. Therefore, we proposed the FD index to quantitatively characterize the pattern of participation of various functional networks across the brain. Through the FD index, we estimated the continuous distribution of functional diversity from the primary cortex to the high-level association cortex. The primary cortex was found to significantly participate in some particular ICN. The high-level association cortex was found to involve in multiple ICNs. The FD index could provide a data-driven estimation about the level of functional diversity, which could be used to detect the change of functional diversity caused by the psychiatric disorders.

## Materials and Methods

In this study, the ICA model was applied to 100 high spatiotemporal resolution rfMRI data from the Human Connectome Project (HCP) (Van Essen et al. 2013). We proposed to estimate the functional diversity both in group-level and in subject-level. Here, we focused on the group-level. More details about the analysis in subject-level would be shown in the supplementary part. There were four steps in the analysis. First, we adopted temporal-concatenated spatial ICA to detect the ICNs based on the preprocessed rfMRI data. Second, we proposed to calculate the index of components homogeneity (CoHo) which reflected the voxel-wise local consistency that a brain region involved with various ICNs. In theory, the voxels around the boundary of two functional networks or the boundary of gray / white matter would show inconsistent patterns of participating in various components, compared with their neighbors. Third, we used the Gaussian Mixture Model (GMM) to estimate the distribution of CoHo values across the brain in order to find the brain regions with high CoHo values. Considering the potential noise and the boundary between functional networks, we restricted the next-step calculation in these brain regions. Finally, we proposed the index of functional diversity (FD index) to further describe the pattern of participation of various ICNs. Fig. 1 gave an overview of the analytic steps.

**Fig 1.**
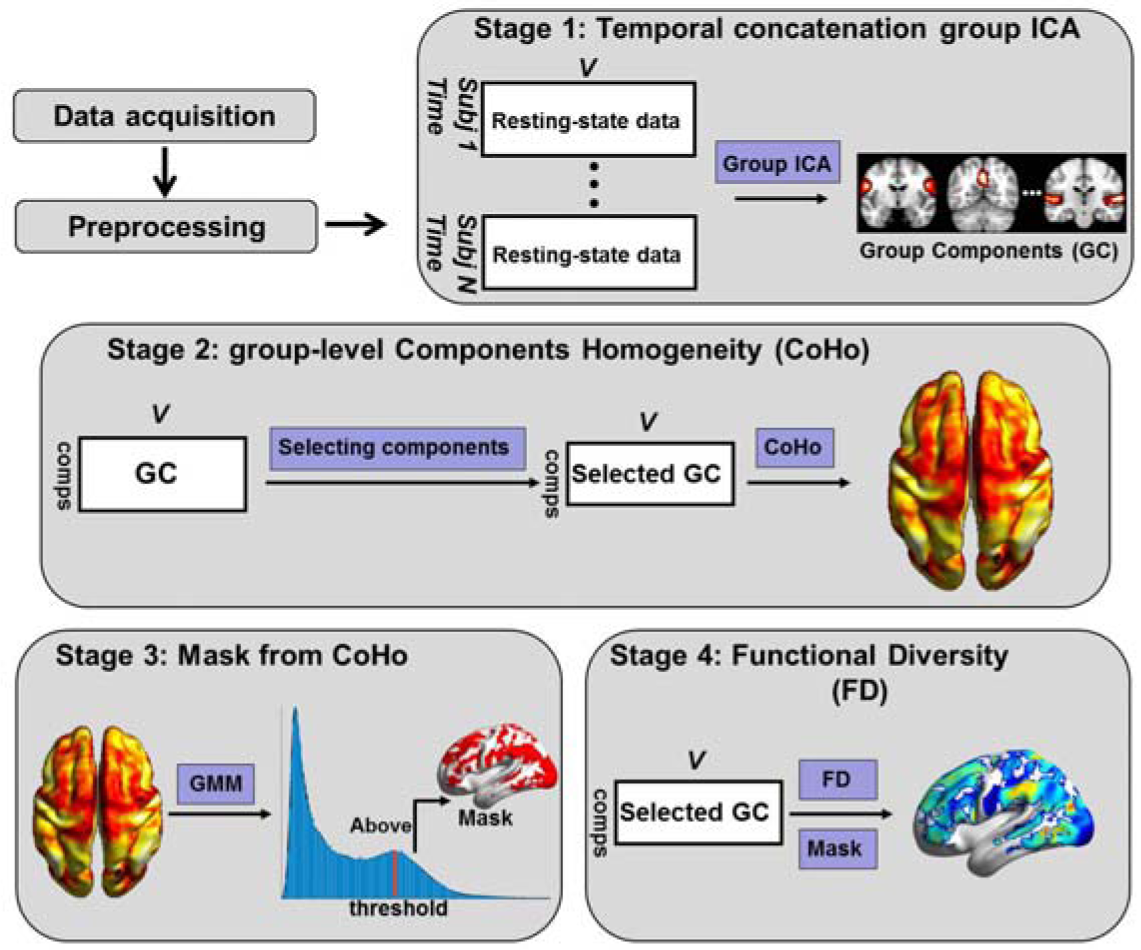
Schematic representation of the analytic steps for calculating the FD index. GMM was the abbreviation of Gaussian Mixture modeling.

### Data acquisition

HCP Q3 data were part of the HCP 500 subjects data released at June 2014 (http://www.humanconnectome.org/data/). There were 100 unrelated subjects in this release. The 100 HCP subjects were between ages 22–35 (mean age was 27.5-31.3; 54 female). As HCP only provided 5-year age range of each participant, the mean age was also estimated as a range. During the resting-state scan, subjects underwent two runs of passive fixation (FIX) in each of two separate sessions. In each session, the phase encoding was in a right to left direction in one run and in a left to right direction in the other run. In this study, we chose the data in session 1 with the left to right phase encoding. Data were acquired on a Siemens Skyra 3 Tesla scanner using a customized SC72 gradient insert and a customized body transmitter coil with 56 cm bore size. Functional data consisted of gradient-echo EPI sensitive to BOLD contrast. Parameters for the resting data were: TR = 720 ms, TE = 33.1 ms, FA = 52°, 2 × 2 × 2 mm voxels, FOV = 208 × 180 mm, and 72 oblique axial slices (Feinberg et al. 2010; Moeller et al. 2010; Setsompop et al. 2012; Van Essen et al. 2012; Yeo et al. 2013). Each functional data contained 1200 time points, i.e. 14.55 minutes.

### MRI image preprocessing

To calculate the functional diversity map, we adopted the preprocessed HCP rfMRI data which were gradient distortion corrected, motion corrected, resampled from the original EPI frames to MNI space and intensity normalized to mean of 10000. More details about the preprocessing step could be found in (Smith et al. 2013). Further, ICA-based artifacts removed data were provided by HCP. In detail, ICA was run in the high-pass filtered preprocessed rfMRI data using FSL’s MELODIC (Beckmann et al. 2005; Beckmann and Smith 2004; Jenkinson et al. 2012; Smith et al. 2004). Then, the decomposed components were fed into FIX (FMRIB’s ICA-based X-noisifier) which had been trained on HCP data as an automated component classifier (Griffanti et al. 2014; Salimi-Khorshidi et al. 2014). The components would be classified as good and bad components. Bad components were then removed from the rfMRI data. For the HCP provided rfMRI data, the cleanup was done in a non-aggressive manner, i.e., only the unique variance associated with the bad components was removed from the data. The 24 motion parameters derived from the motion estimation were also regressed out of the data (Satterthwaite et al. 2013). After these steps, the ICA-FIX denoised rfMRI volumetric time-series were provided.

To determine the functional connectivity pattern of brain regions with different level of functional diversity, we further processed the ICA-FIX denoised rfMRI data using FMRIB Software Library (FSL: version 5.0) and Analysis of Functional NeuroImages (AFNI). A series of preprocessing steps were performed: (1) band-pass filtering the time-series (0.01 Hz < f < 0.1 Hz); (2) regressing the nuisance signals including white matter mean signal, cerebrospinal fluid mean signal and global mean signal; (3) spatial smoothing the residuals using 4-mm full-width at half-maximum (FWHM) Gaussian kernel. Here, the resting-state functional connectivity was based on the Pearson correlation calculated on the processed rfMRI data. For each subject, the correlation coefficients between the mean time series of specified brain region and that of each voxel of the whole brain were calculated. These coefficients were converted to Z-values using Fisher’s Z transformation. A one-sample t-test was adopted to determine the brain regions significantly correlating with the specified brain region in a voxel-wise manner. Type I errors in multiple comparisons were controlled by the false discovery rate (FDR).

### Temporal concatenation group ICA

In the group level, Probabilistic Independent Component Analysis (PICA) was adopted to decompose the temporally concatenated data of individual subject into different spatial components (Fig. 1: Stage 1) (Beckmann et al. 2005; Beckmann and Smith 2004). PICA decomposed the rfMRI dataset into a linear mixture of spatially independent components plus Gaussian noise. Probabilistic principal component analysis (PPCA) was used to infer a set of spatially whitened observations. Then, the noise covariance structure could be estimated from the residuals after PPCA. The noise was used for transforming the raw ICs to Z-statistic maps that represented the amount of variability explained by the entire decomposition at each voxel location relative to the noise (Beckmann et al. 2005). The high Z-statistic value illustrated the low probability that the signal in this voxel was generated by noise. That is, any independent information would have to reveal itself via its derivation from Gaussianity (Beckmann et al. 2005). In addition, the ICA decomposition was obtained using FastICA scheme to optimize for non-Gaussianity source estimates (Hyvarinen 1999).

In the current study, we applied the temporally concatenated PICA via Melodic (Melodic 3.13). The number of independent components was set as 20. The 20-components were considered as the lower order that represented large-scale ICNs (Smith et al. 2009). We also tried different number of components (30, 40, 50, 60 and 70) to evaluate the functional diversity map across various orders of ICA. In addition, it was considered as high model order to set the number of component as 70 that could result in more sub-networks (Allen et al. 2011; Kiviniemi et al. 2009).

### ICN selection

Before the next step analysis, the components from ICA decomposition should be classified as corresponding ICNs and artificial non-ICNs. The up-threshold Z-score maps (|Z-score|>5) of components were used as input for the classification. The components classified as ICNs had the characteristics: 1) focused on cortical structures; 2) clustered voxel groups. Meanwhile, the components considered as artificial non-ICNs had the features: 1) located at the borders of the brain; 2) located in cerebrospinal fluid; 3) located at the proximity of ventricle; 4) located at white matter; 5) dispersedly distributed around the brain without large clustered voxel. Two viewers (Congying & Lingzhong) selected the components following the described criteria separately. Only components confidently classified as ICNs were preserved for the next-step analysis.

### Component Homogeneity (CoHo)

Before considering the functional diversity across the brain, we firstly defined the CoHo index to measure the local consistency, i.e., the consistent level of brain regions to participate across various ICNs. We thought that the functional diversity would reflect underlying functional organization only in brain regions with high CoHo values. Brain regions with low CoHo values might reflect the instability of the corresponding signals, such as the effect of noise. In detail, the CoHo index was defined as:

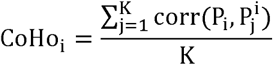

where CoHo_*i*_ was the component homogeneity at voxel i (Fig. 1: Stage 2). K was the neighbor voxels around voxel i. K could be set as 6, 18 and 26 corresponding to different definition of neighbor. Here, K was set as 26 for the nearest neighbor in three-dimensional space. P_*i*_ was the vector composed of Z-scores across the selected ICNs at voxel i. 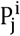 had the same definition of P_*i*_, which was jth neighbor of voxel i. corr *,* was the Pearson correlation coefficient between two vectors. Then, CoHo_*i*_ was transformed as Z_CoHo__i__ using Fisher’s z-transform, which is:

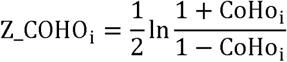

where ln(*) was the natural logarithm. Here, the transformation was used to extend the scale of CoHo_*i*_ for clearly showing the various level of local consistency around the cortex.

### Gaussian Mixture Model (GMM)

Inspired by Melodic in which GMM was firstly adopted to inference the activation part in each component (Beckmann and Smith 2004), we also modeled the distribution of CoHo values across the brain via GMM. To be noticed, as an alternative to GMM, Melodic had been changed to use one Gaussian distribution and two Gama distribution to model the Z-scores in each component referring to (Hartvig and Jensen 2000). Here, we focused on GMM which had the form as:

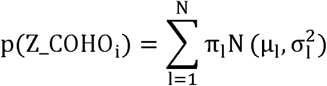

where π_1_ was the mixture coefficient for the lth Gaussian distribution. 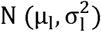 was a Gaussian distribution with mean of µ_1_ and variance of 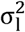. N was the number of Gaussian distribution. p Z_COHO was the probability density at the value Z_COHO_*i*_ which had the same mean as described above. Using a sufficient number of Gaussian distribution, GMM could model the probability density function of any input signal (Bishop 2006). Therefore, an analytical expression of the distribution of CoHo value could be deduced. The estimation of the parameters for the model was based on the expectation-maximization (EM) algorithm (Bishop 2006). The number of mixed Gaussian distribution was decided by the Bayesian information criterion (Schwarz 1978). We thought that the deduced function of mixture distribution would have peaks and valleys that reflected the underling structure of ICNs. Some peak could be used as a threshold to generate the brain regions with high values.

### Functional diversity (FD)

After calculating the CoHo values across the brain, we defined the index of functional diversity as:

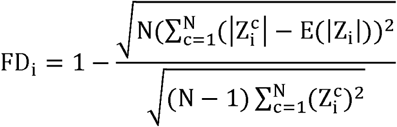

where 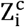 was the Z-scores at the voxel i within component c. N was the number of the selected components. FD_*i*_ ∈ [0,1] was the index of functional diversity at voxel i. More specially, FD was an index that measured the level of anisotropic distribution related with participation of various components. If a voxel equably participated in each separate ICN, FD value at this voxel would be calculated as 1. Conversely, if the voxel specifically participated in only one ICN, FD value would be 0. To be notice, FD value would be high at some voxel with random Z-scores, such as the voxels located in white matter. Therefore, we emphasized to calculate FD at brain regions with high CoHo values. These regions were related with significant signal expression of some ICNs, which were impossible to be mainly generated by noise or uncertainties in calculating PICA model. In addition, we adopted the absolute Z-scores for calculating FD inspired by the inference process in Melodic. By using Gaussian and Gama distribution, the inference of significant activation or deactivation was simultaneously modeled as probability in Melodic. The significant activation or deactivation in some component was related with the derivation from the Gaussian noise. The further the derivation, the more possible the activation/deactivation obtained information. Therefore, we used the absolute Z-score to model the extent that a voxel was involved with a particular component. In other words, we only considered the extent about participation of ICN without consideration of the positive or negative pattern. After masking the derived functional diversity map, we spatially smoothed the masked map using 4-mm FWHM Gaussian kernel. To be notice, the mask adopted here only included the cortical structure. The subcortical tissues were excluded by using the max-probability atlases with 0.25 probability included in FSL software (Harvard-Oxford subcortical structural atlases kindly provided by the Harvard Center for Morphometric Analysis). The cerebellum was excluded based on the probabilistic MR atlas of the human cerebellum with 0.25 probability (Diedrichsen et al. 2009).

## Results

We here focused on the group-level results based on the group ICA decomposition of 20 components. The results based on subject-level or more model orders were shown in the supplementary part. In addition, there were some abbreviations for the names of brain regions used in the following part: temporoparietal junction (TPJ), parietal-temporal-occipital junction (PTOJ), temporal-occipital junction (TOJ), anterior cingulate cortex (ACC), middle cingulate cortex (MCC), posterior cingulate cortex (PCC), middle frontal gyrus (MFG), superior frontal gyrus (SFG), inferior parietal lobule (IPL), superior parietal lobule (SPL), middle temporal gyrus (MTG), dorsolateral frontal cortex (DLPFC), middle occipital gyrus (MOG), inferior frontal gyrus (IFG).

### CoHo map and the GMM estimation

Two viewers (Congying & Lingzhong) selected 19 components from 20 decomposed components as the reflection of underlying ICNs. The non-selected component was related with signal from white matter. The selected components were shown in Supplemental Fig. 1. Figure 2 presented the distribution of group-level CoHo and the deduced brain regions with high CoHo values. The group-level CoHo map without threshold was shown in Fig. 2A &2B as surface view and slices separately. Illustratively, regions with evidently high CoHo value from Fig. 2A mainly included visual cortex (significantly distributed in visual area V1 & V2) and primary sensory-motor cortices. As the related signal had been selected out, the CoHo values of white matter were significantly low (Fig. 2B). There still were some brain regions with relatively high CoHo values comparing with ones of white matter. These regions mainly distributed in association cortex of TPJ and PTOJ, PCC, precuneus and MFG.

**Fig 2.**
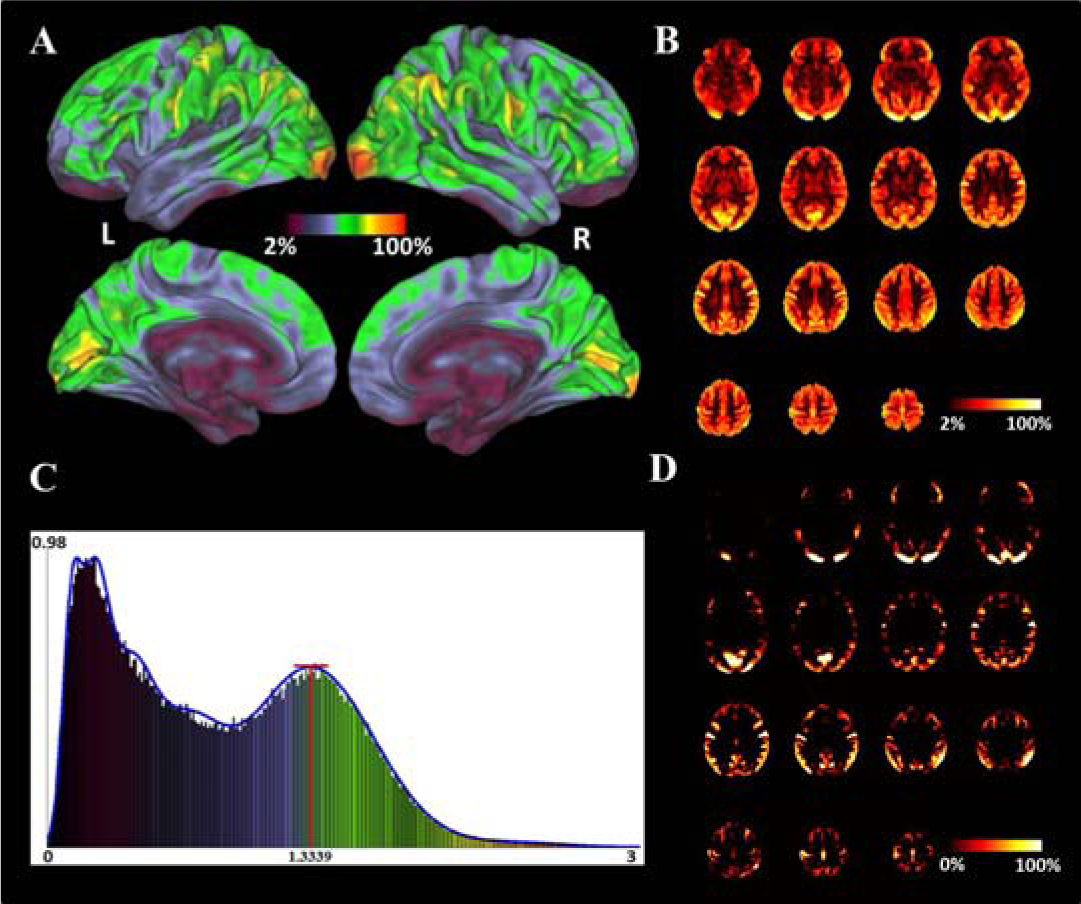
The group-level CoHo map and the GMM estimation. (A) The group-level CoHo map projected on surface. (B) The transverse slices of group-level CoHo map. (C) The estimation of GMM. The blue curve was the Gaussian mixture function to model the underlying probability density function. The chromatic histogram was derived from the CoHo map. The color here was corresponding to the color on Fig.2A. The red line was the chosen threshold. (D) The transverse slices of the brain regions with CoHo values greater than the threshold.

The estimation of GMM on the calculated CoHo map was shown in Fig. 2C. According to the Bayesian information criterion, the number of mixed Gaussian distributions was chosen as 8. The mixture of these eight Gaussian distributions was shown in Fig. 2C. There were obvious two peaks and one valley in the estimated density function. We found that the existence of valley was resulted from the different CoHo values between white matter and grey matter (Fig. 2A & 2C). In order to define the next-step brain mask with more confidence, we chose a relatively high threshold that was the CoHo value corresponding to the last peak location (the red line in Fig. 2C). The location was detected based on the derivative of the estimated probability density function. The brain regions passing through the threshold were shown in Fig. 2D as slices. We found that the white matter was significantly excluded.

### Functional diversity (FD)

We calculated the defined FD index. The results were shown in Fig. 3. As an overview, the FD index within the derived mask was shown in Fig. 3A. We found that the primary cortex mainly including visual cortex and primary sensory and motor cortices had low FD values. The association cortices and higher order association cortices presented high FD values, for example, precuneus, IPL and TOJ. In order to find the illustrative brain regions with sufficient low or high FD values, we checked the histogram of the FD values within the mask (Fig. 3B). We separately chose the brain regions with FD values of top or bottom 5%. We further adopted a cluster-size correction for the derived brain regions, i.e., only the brain regions with cluster-size greater than 100 voxels were reserved. The corrected results were shown in Fig. 3C. The brain regions with significant low FD values were distributed in visual area V1 & V2 and bilateral postcentral gyrus (left side in Fig. 3C). Meanwhile, there were brain regions with significant high FD values, which included precuneus, PCC, SFG, left supramarginal gyrus, left IPL, left MTG and left TOJ (right side in Fig. 3C).

**Fig 3.**
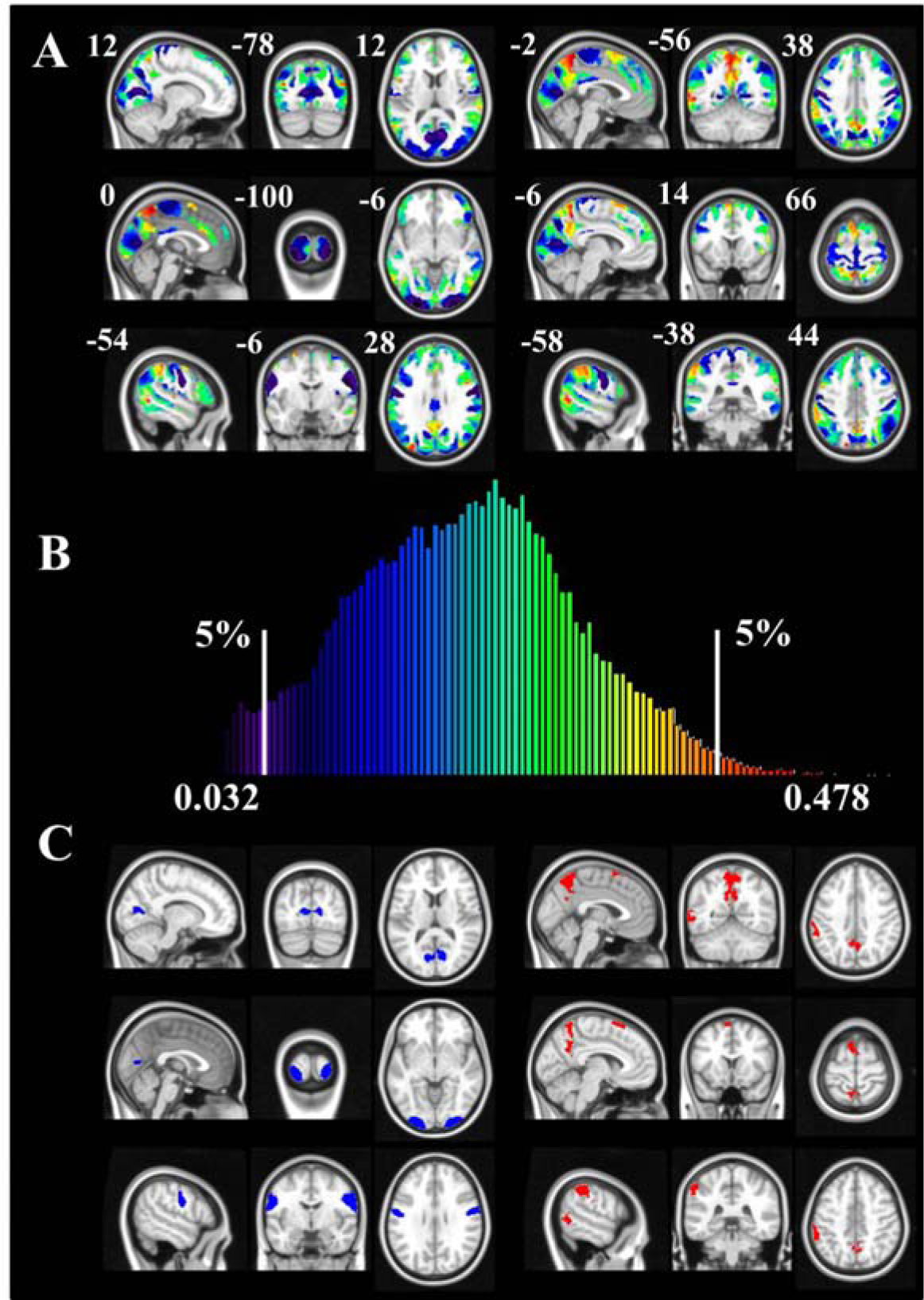
The distribution of FD index. (A) The derived FD index within mask. (B) The histogram of FD index. The color in the histogram was corresponding to the color on Fig.3A. (C) The significant brain regions with low or high FD values. The blue regions in left side were corresponding to low FD values. The red regions in right side were corresponding to high FD values.

### Participation of ICNs and functional connectivity

In order to illustrate the underling structures resulting in different FD values, we analyzed the patterns of participation of ICNs in the brain regions with significant low or high FD values (just as described above). We adopted probability map of each components generated by fitting the model of one Gaussian and two Gama distributions. Then, the participation of ICNs with each brain region was defined as the mean probability of this region participating in the corresponding component.

There were five derived regions with low FD values, which were named as ROI1 to ROI5 (surface view saw in Fig. 4). We showed the fingerprints that were the mean probabilities participating in different ICNs. We noticed the common feature in these ROIs, which was the significant participation (mean probability > 0.5, the light grey circle in Fig.4 corresponding to the probability of 0.5) only involved with some particular ICN. More specially, ROI1 & ROI2 which were mainly distributed in left /right visual area V1 and V2 were significant participated in independent component 3 (IC3; the red and yellow ROIs in Fig. 4). All these independent components could be seen in Supplemental Fig. 1. From the distribution of IC3, we found that up-threshold part of IC3 was mainly shown up in the central-field representation of left /right visual areas V1 and V2. ROI3 located in primary visual cortex (V1), which specially showed in IC2 (the green ROI in Fig. 4). IC2 mainly occupied peripheral V1 and V2. ROI4 & ROI5 were mainly shown up in bilateral postcentral gyrus, which were significantly related with IC14 (the blue and purple ROIs in Fig. 4). The up-threshold part of IC14 was mainly located in bilateral postcentral gyrus. We found that the ICs related with these ROIs almost distributed in special and localized brain areas.

**Fig 4.**
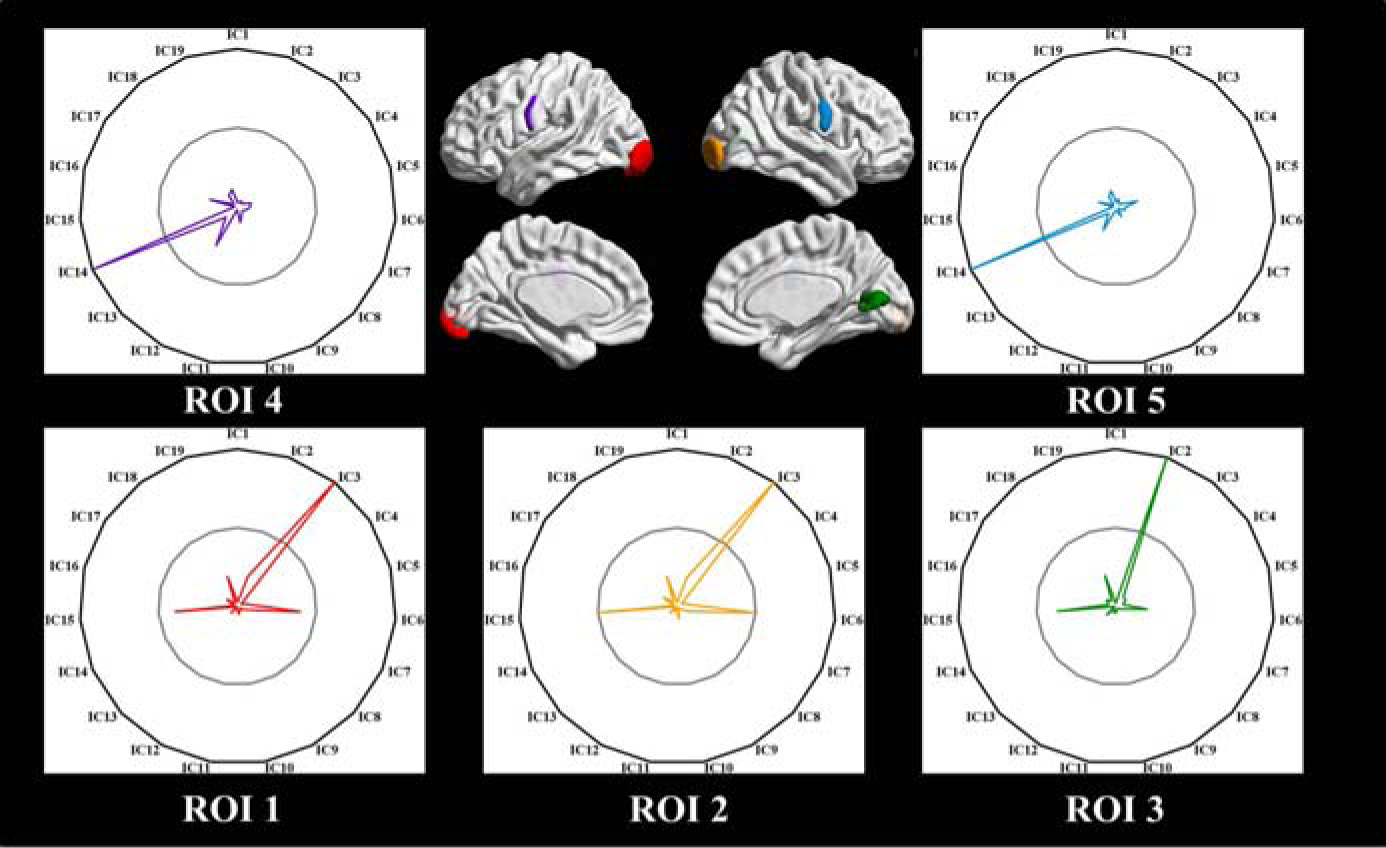
The pattern of ICNs-participation for the derived brain regions with low FD values. The color of the brain region on the surface was corresponding with the color in the fingerprint. We named the brain regions as ROI1 (red), ROI2 (yellow), ROI3 (green), ROI4 (purple) and ROI5 (blue). The grey circle in the fingerprint represented the probability of 0.5. The black circle represented the probability of 1.

We further calculated the resting-state functional connectivity for these five ROIs to provide more insights for the properties of these ROIs (Fig. 5). The threshold used for testing the significance of functional connectivity was set as p < 0.001 (FDR corrected) and cluster-size > 200 voxels. ROI1 & ROI2 showed positive correlation with each other, and negative correlation with somatosensory and somatomotor cortex. ROI3 was positively correlated with bilateral V1, and negatively correlated with the bilateral supramarginal cortex. ROI4 & ROI5 positively correlated with each other. Some parts of DLPFC were found to negatively correlate with ROI4 & ROI5.

**Fig 5.**
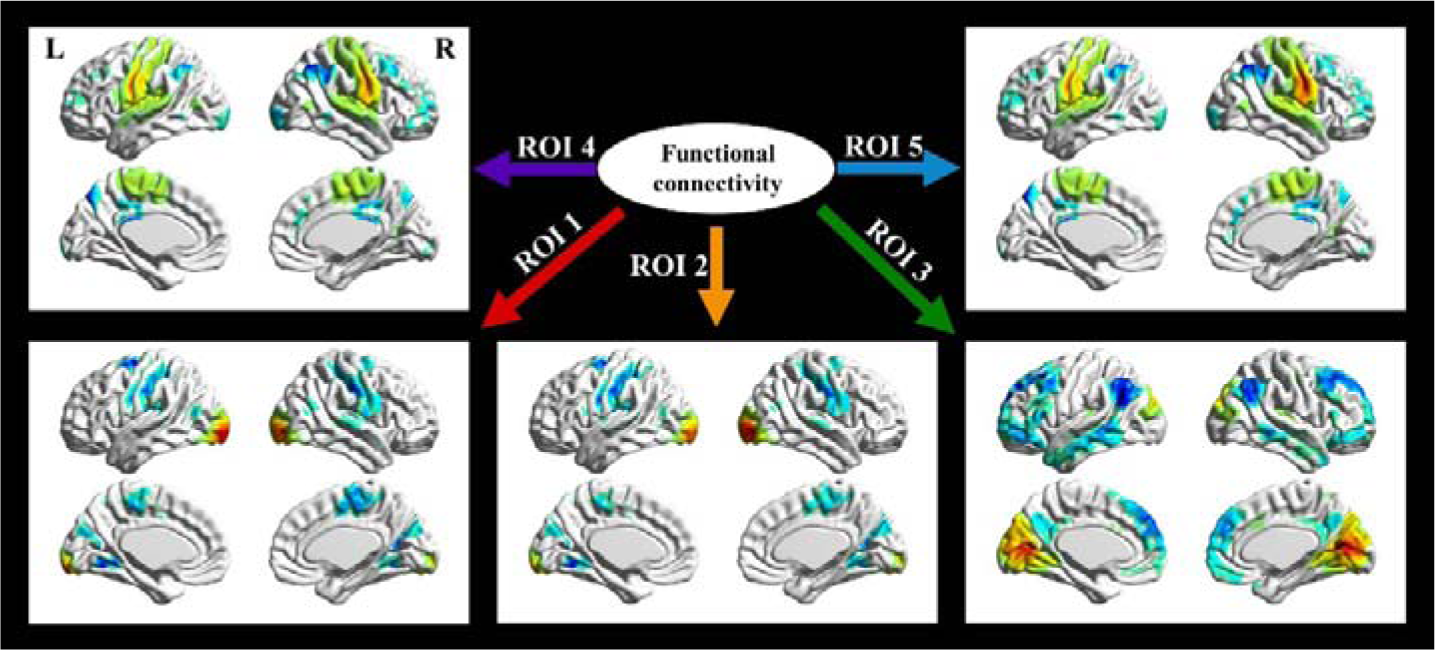
The pattern of functional connectivity for the five ROIs with low FD value. One sample t-test was adopted to test the significance of the functional connectivity with FDR-correction at p < 0.001 and cluster-size > 200 voxels.

There still were five derived brain regions with high FD values. We used ROI 6 to ROI 10 to name these brain regions. We gave the surface view and the finger-prints of the ICNs-participation of these ROIs (Fig. 6). We found that the brain regions with high FD values presented complex patterns of participation of ICNs. These regions significantly participated in various ICNs. In detail, ROI6, which mainly located in precuneus and PCC, participated in IC1, 4, 7, 8, 10, 11 and 13 (the yellow ROI in Fig. 6). ROI7 was mainly shown up in left IPL, which participated in IC5, 6, 7, 8, 10, 13, 18 and 19 (the green ROI in Fig. 6). ROI8 was found in SFG, which had the participation in IC1, 5, 7, 9, 13, 18 and 19 (the blue ROI in Fig. 6). ROI9 was shown up in left MTG, which participated in IC5, 6, 7, 8, 9, 10 and 19 (the red ROI in Fig. 6). ROI10 located in left TOJ, which participated in IC1, 4, 6, 10, 12, 15, 18 and 19 (the orange ROI in Fig. 6). The related ICs were shown in Supplemental Fig. 1. We found that the ICNs related with these ROIs were pervasively distributed, rather focused on some localized regions.

**Fig 6.**
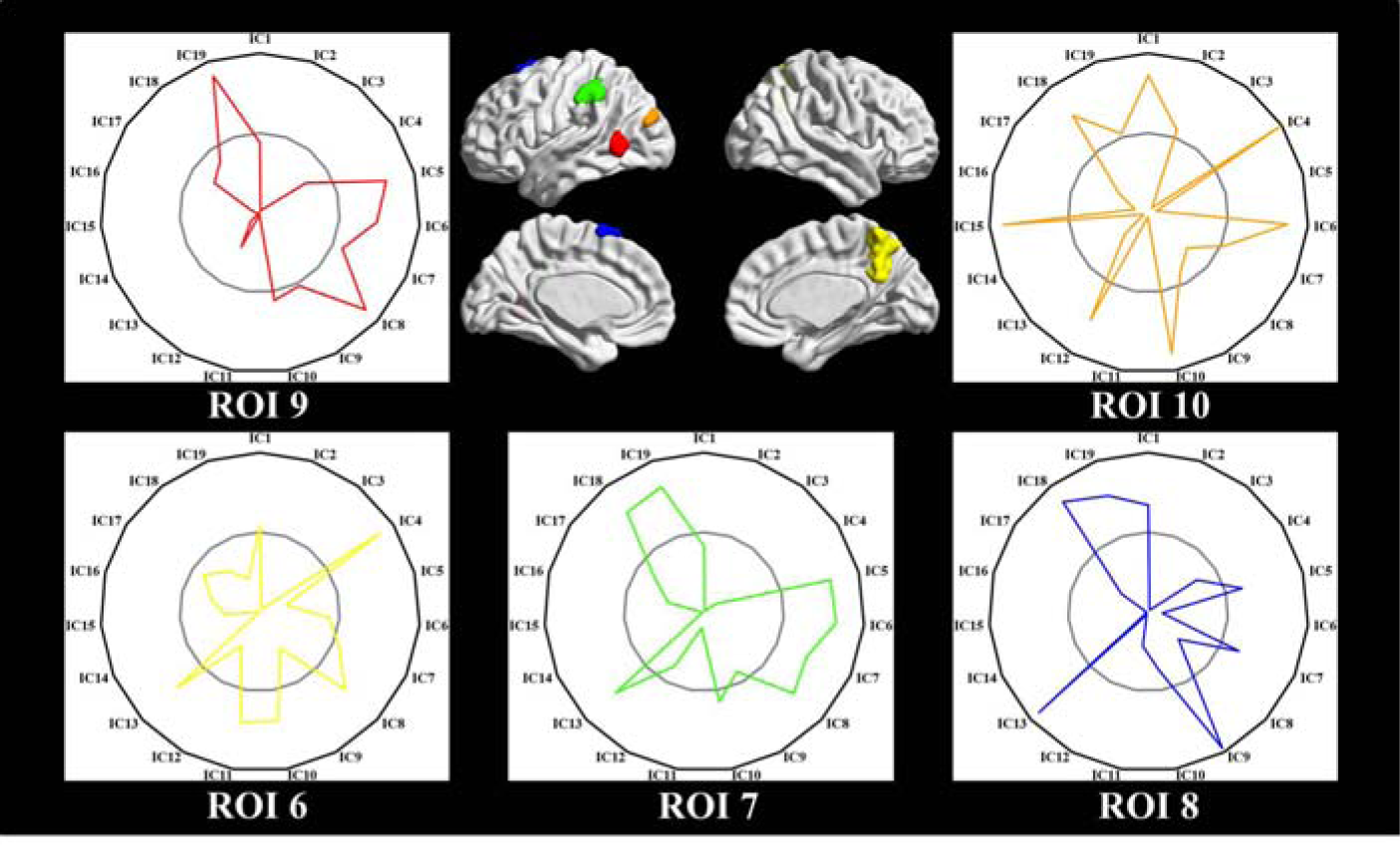
The pattern of ICNs-participation for the derived brain regions with high FD values. The color of the brain region on the surface was corresponding with the color in the fingerprint. We named the brain regions as ROI6 (yellow), ROI7 (green), ROI8 (blue), ROI9 (red) and ROI10 (orange). The grey circle in the fingerprint represented the probability of 0.5. The black circle represented the probability of 1.

The pattern of functional connectivity was recognized for these five ROIs with high FD value. The maps of functional connectivity were shown in Fig. 7, which were corrected on FDR (p < 0.001) and cluster-size (> 200 voxels). More specially, ROI6 presented positive functional connectivity with its contralateral part, bilateral angular gyrus and MFG. The negative correlation was shown up in somatosensory and somatomotor cortex and insular. ROI7 positively correlated with its contralateral part, some part of DLPFC, insular, MTG and SFG, and, negatively correlated with precuneus, PCC, MFG and bilateral MOG. ROI8 showed positive correlation with SFG, MFG, angular gyrus and some part of DLPFC, and, negative correlation with precuneus and MOG. ROI9 had positive functional connectivity with its contralateral region, bilateral IPL and left IFG. The negative functional connectivity was found in primary visual area, PCC, and MFG. ROI10 showed positive correlation with its contralateral part, visual cortex, SPL, and precuneus, and, negative correlation with MFG, ACC, MCC and bilateral angular gyrus.

**Fig 7.**
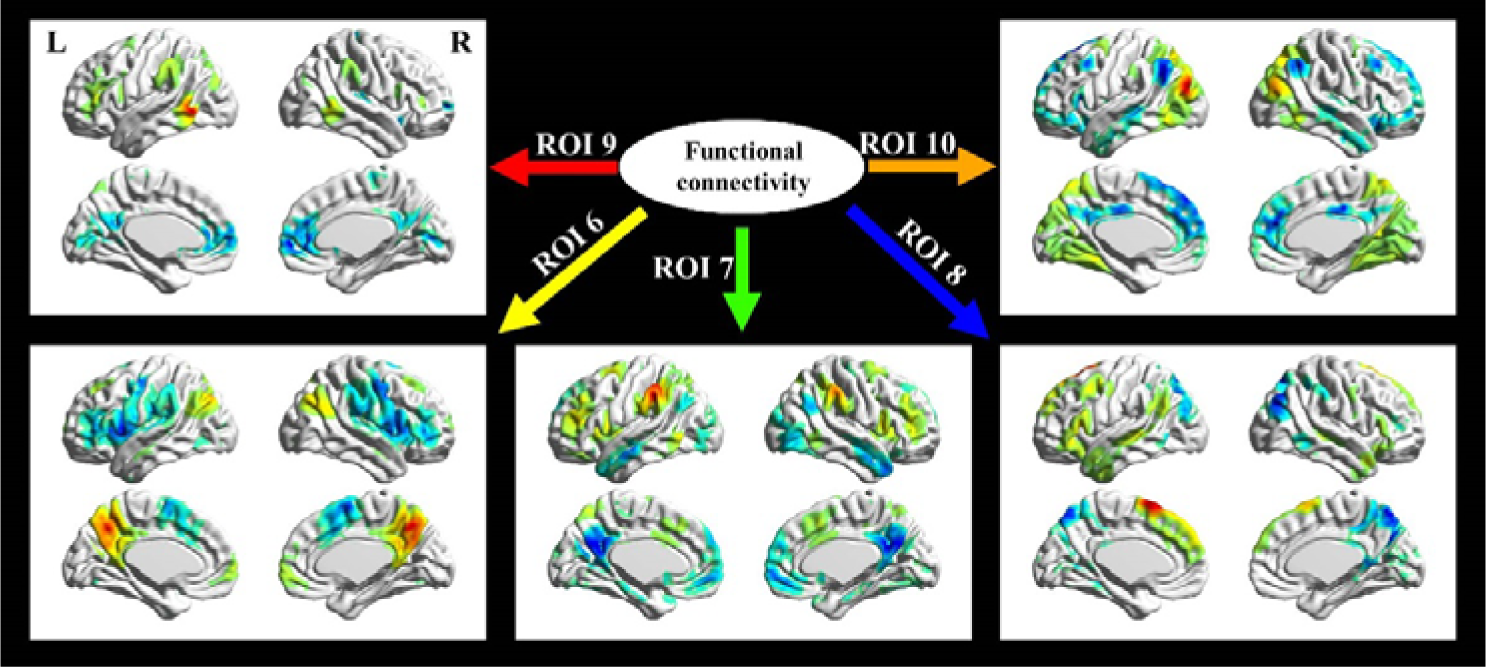
The pattern of functional connectivity for the five ROIs with high FD values. One sample t-test was adopted to test the significance of the functional connectivity with FDR-correction at p < 0.001 and cluster-size > 200 voxels.

## Discussion

In the present study, we utilized the voxel-wise participation of ICNs detected by sICA to estimate the functional diversity across brain. The CoHo index was firstly defined to decide the brain regions with homogeneous pattern of participating in various ICNs. Then, the FD index was calculated within the brain regions decided by CoHo index in a voxel-wise manner. We further illustrated the detailed fingerprints of participant patterns found in brain regions with high or low FD values to explore the cause of different FD values.

The hypothesis that the organization of human brain function depends on the network structure among different brain regions is supported by a substantial body of evidence (Bressler 1995; Bressler and Menon 2010; M. Mesulam 2009; M. M. Mesulam 1990; Rubinov and Sporns 2010; Sporns 2013, 2014). The segregation and integration between brain regions, such as the modular structure and the rich-club organization, have been estimated by using network model (Power et al. 2011; Sporns 2013; van den Heuvel and Sporns 2011; Yeo et al. 2011). Based on the hypothesis of network, the functional properties of brain regions, such as functional connectivity and functional co-activation pattern, within a special network should be more similar than the brain regions located in different networks. In the perspective of clustering, brain regions attributed to a special network could be clustered as one cluster using the functional or structural features (Fan et al. 2014; Yeo et al. 2011). The functions related with the specialized network have been estimated by recent studies (Laird et al. 2011; Smith et al. 2009). Especially, the resting-state networks are similar with ‘intrinsic’ functional network architecture presenting across many tasks (Braga et al. 2013; Cole et al. 2014). The full repertoire of functional networks utilized in action of the brain is found to continuously show up at resting-state (Smith et al. 2009). So, the function-specialized networks could be found in resting-state. Based on network perspective, new properties, especially in large scale, have been found in the functional organization of human brain. For example, Power et al. has proposed to define the hubs in human brain using the network as object of study, which are different with the voxel-wise defined hubs (Power et al. 2013). Braga et al. has estimated the echoes of brain by using a searchlight approach over the whole brain to find the particular searchlights involved with multiple functional networks (Braga et al. 2013). These found searchlights were believed to contain the echoes of the neural signals from multiple functional networks. It was supposed that these found searchlights played important roles in connecting various functional networks. The network perspective maybe opens a new avenue to further understand the function properties of human brain, especially in large scale (Pessoa 2014). We also adopted the network perspective in our study to describe the large-scale functional diversity of brain regions to some extent.

The backbone of our study was the detection of the network structures in human brain. As we focused on the data from rfMRI, seed-based functional connectivity and ICA-based decomposition had emerged as two dominant approaches commonly used to detect ICNs (Biswal et al. 1997; De Luca et al. 2006). The ICA-based identification of ICNs had been characterized with high robustness and test-retest reliability (Damoiseaux et al. 2006; Zuo et al. 2010). The estimation of ICNs under a resting condition had been described as a ‘killer application’ of ICA (Beckmann 2012). More specially, we adopted the PICA to estimate the ICNs. The statistically independent source signal was hypothesized to compose of non-Gaussian signal plus Gaussian noise. Using the process of normalization of voxel-wise residual noise, the variances of Gaussian noises in different source signals were normalized as unit variance. The Gaussian/Gamma mixture approach modeled the normalized Gaussian distribution for the background noise and two Gamma distributions for the activations and de-activations. So, the voxel-wise Z-statistics in normalized independent component maps illustrated the extent that the activation in corresponding voxel was affected by normalized Gaussian noise. For instance, a high Z-statistic value at a voxel illustrated that the activation at the voxel was barely generated by the Gaussian noise. It had been proposed that the voxel-wise Z-statistics could be used to illustrate the extent that the voxel was significantly modulated by the time courses of independent component (Beckmann 2012). Therefore, we proposed to use the voxel-wise Z-statistic to measure the extent that a voxel participated in particular ICN.

CoHo index here was proposed to detect the brain regions which consistently participated in various ICNs. The index was defined in voxel level, which concerned the consistency between a voxel and its neighboring voxels. The voxel with low regional consistency was suspected to be affected by the noise of the data or boundary effects, i.e., located in the boundary of various ICNs. For example, we had observed that the voxels in temporal lobe and in orbitofrontal cortex had low CoHo values (Fig. 2A). Large bold signal loss in these regions had been found in other study (Wig et al. 2014). So, we suspected that the low CoHo value here was caused by the signal loss. The brain regions with high CoHo values would attract more interests for the next-step calculation of functional diversity, as these regions reflected the functional properties with more confidence in the current dataset. Attempting to detect the brain regions with high CoHo values, we adopted the GMM model to calculate the probability density distribution of the CoHo value. We observed a bimodal pattern in the distribution (Fig. 2C). The first peak was corresponding to the distribution of CoHo value in white matter. The second peak was related with the CoHo value in gray matter. Through CoHo index, we could observe the different properties between white matter and gray matter. We used the CoHo value corresponding to the second peak as the threshold to define the brain regions with high CoHo values. In fact, the approach based on threshold was somewhat arbitrary. We considered this threshold as a strict one, as some part of gray matter was excluded according to the threshold. We tended to adopt a conservative threshold to detect the brain regions with consistent participant pattern of ICNs.

The functional diversity was described by FD index that was defined as the extent of the isotropic participation of ICNs. If a brain region participated in various ICNs evenly, this region would have a high FD value. The brain regions with non-significant participation of ICNs, such as part of white matter, would have a high FD value, as the Z-statistics in these regions were mainly affected by Gaussian noise. Therefore, the mask from CoHo map was adopted to preserve that the FD value was not mainly caused by noise. We observed that the primary cortex showed lower FD values, such as the sensory and motor cortices. Some part of association cortex showed much higher FD values. From the selected brain regions that had significantly low or high FD values, the primary cortex with low FD values only significantly participated in one ICNs (Fig. 4). The association cortex with high FD values represented a more complex pattern of participation, which significantly participated in various ICNs. ICNs detected in resting-state had correspondence with the co-activation structures found in task state (Smith et al. 2009). These ICNs are supposed to obtain functional specialization. In large scale, the brain region with low FD value, i.e., the participation of unique ICN, might illustrate that the region was related with unique function. On the other hand, the region involved in various brain functions might participate in several ICNs. The functional connectivity of these selected regions further validated the difference function properties between brain regions with low / high FD values. We found that the regions with low FD values mainly had significant functional connectivity with their contralateral parts. The brain regions with high FD values showed a more distributed pattern of functional connectivity. It was corresponding with other findings that the higher-order brain systems obtained long-range functional connectivity (Sepulcre et al. 2012; Tomasi and Volkow 2010, 2011).

In summary, we adopted the perspective of networks to study the functional properties of human brain in large scale. Our approach allowed us to quantitatively characterize the degree of the participation of ICNs across the brain by FD index. FD index could be considered as a potential index to characterize the functional diversity. It was noticed that the functional diversity changed from primary cortex to high-order association regions revealed by a quantitative index. In future, a particular interesting application of the FD index was involved with the application to the psychiatric disorders, such as schizophrenia which was suspected to be related with disabled integration of various functions.

## Supporting information

## Acknowledgements

This work was partially supported by the National Key Basic Research and Development Program (973) (Grant No. 2011CB707800) and the Strategic Priority Research Program of the Chinese Academy of Sciences (Grant No. XDB02030300). Data were provided by the Human Connectome Project, WU-Minn Consortium (Principal Investigators: David Van Essen and Kamil Ugurbil; 1U54MH091657) funded by the 16 NIH Institutes and Centers that support the NIH Blueprint for Neuroscience Research; and by the McDonnell Center for Systems Neuroscience at Washington University.

## References

Allen, E. A., Erhardt, E. B., Damaraju, E., Gruner, W., Segall, J. M., Silva, R. F., et al. (2011). A baseline for the multivariate comparison of resting-state networks. Frontiers in systems neuroscience, 5, 2.

Anderson, M. L., Kinnison, J., & Pessoa, L. (2013). Describing functional diversity of brain regions and brain networks. Neuroimage, 73, 50–58, doi:10.1016/j.neuroimage.2013.01.071.

Beckmann, C. F. (2012). Modelling with independent components. Neuroimage, 62(2), 891–901, doi:10.1016/j.neuroimage.2012.02.020.

Beckmann, C. F., DeLuca, M., Devlin, J. T., & Smith, S. M. (2005). Investigations into resting-state connectivity using independent component analysis. Philosophical Transactions of the Royal Society B-Biological Sciences, 360(1457), 1001–1013, doi:DOI 10.1098/rstb.2005.1634.

Beckmann, C. F., & Smith, S. A. (2004). Probabilistic independent component analysis for functional magnetic resonance imaging. Ieee Transactions on Medical Imaging, 23(2), 137–152, doi:DOI 10.1109/Tmi.2003.822821.

Bishop, C. M. (2006). Pattern recognition and machine learning (Information science and statistics). New York: Springer.

Biswal, B. B., Van Kylen, J., & Hyde, J. S. (1997). Simultaneous assessment of flow and BOLD signals in resting-state functional connectivity maps. NMR Biomed, 10(4-5), 165–170.

Braga, R. M., Sharp, D. J., Leeson, C., Wise, R. J. S., & Leech, R. (2013). Echoes of the Brain within Default Mode, Association, and Heteromodal Cortices. Journal of Neuroscience, 33(35), 14031–14039, doi:DOI 10.1523/Jneurosci.0570-13.2013.

Bressler, S. L. (1995). Large-scale cortical networks and cognition. Brain Res Brain Res Rev, 20(3), 288–304.

Bressler, S. L., & Menon, V. (2010). Large-scale brain networks in cognition: emerging methods and principles. Trends Cogn Sci, 14(6), 277–290, doi:10.1016/j.tics.2010.04.004.

Cole, M. W., Bassett, D. S., Power, J. D., Braver, T. S., & Petersen, S. E. (2014). Intrinsic and task-evoked network architectures of the human brain. Neuron, 83(1), 238–251, doi:10.1016/j.neuron.2014.05.014.

Damoiseaux, J. S., Rombouts, S. A., Barkhof, F., Scheltens, P., Stam, C. J., Smith, S. M., et al. (2006). Consistent resting-state networks across healthy subjects. Proc Natl Acad Sci U S A, 103(37), 13848–13853, doi:10.1073/pnas.0601417103.

De Luca, M., Beckmann, C. F., De Stefano, N., Matthews, P. M., & Smith, S. M. (2006). fMRI resting state networks define distinct modes of long-distance interactions in the human brain. Neuroimage, 29(4), 1359–1367, doi:10.1016/j.neuroimage.2005.08.035.

Diedrichsen, J., Balsters, J. H., Flavell, J., Cussans, E., & Ramnani, N. (2009). A probabilistic MR atlas of the human cerebellum. Neuroimage, 46(1), 39–46, doi:10.1016/j.neuroimage.2009.01.045.

Fan, L., Wang, J., Zhang, Y., Han, W., Yu, C., & Jiang, T. (2014). Connectivity-based parcellation of the human temporal pole using diffusion tensor imaging. Cerebral Cortex, 24(12), 3365–3378, doi:10.1093/cercor/bht196.

Feinberg, D. A., Moeller, S., Smith, S. M., Auerbach, E., Ramanna, S., Gunther, M., et al. (2010). Multiplexed echo planar imaging for sub-second whole brain FMRI and fast diffusion imaging. PLoS One, 5(12), e15710, doi:10.1371/journal.pone.0015710.

Felleman, D. J., & Van Essen, D. C. (1991). Distributed hierarchical processing in the primate cerebral cortex. Cerebral Cortex, 1(1), 1–47.

Fox, P. T., Laird, A. R., Fox, S. P., Fox, P. M., Uecker, A. M., Crank, M., et al. (2005). BrainMap taxonomy of experimental design: description and evaluation. Hum Brain Mapp, 25(1), 185–198, doi:10.1002/hbm.20141.

Fox, P. T., & Lancaster, J. L. (2002). Opinion: Mapping context and content: the BrainMap model. Nat Rev Neurosci, 3(4), 319–321, doi:10.1038/nrn789.

Goldmanrakic, P. S. (1988). Topography of Cognition - Parallel Distributed Networks in Primate Association Cortex. Annual Review of Neuroscience, 11, 137–156, doi:DOI 10.1146/annurev.ne.11.030188.001033.

Griffanti, L., Salimi-Khorshidi, G., Beckmann, C. F., Auerbach, E. J., Douaud, G., Sexton, C. E., et al. (2014). ICA-based artefact removal and accelerated fMRI acquisition for improved resting state network imaging. Neuroimage, 95, 232–247, doi:10.1016/j.neuroimage.2014.03.034.

Hartvig, N. V., & Jensen, J. L. (2000). Spatial mixture modeling of fMRI data. Hum Brain Mapp, 11(4), 233–248.

Hyvarinen, A. (1999). Fast and robust fixed-point algorithms for independent component analysis. Ieee Transactions on Neural Networks, 10(3), 626–634, doi:DOI 10.1109/72.761722.

Jenkinson, M., Beckmann, C. F., Behrens, T. E., Woolrich, M. W., & Smith, S. M. (2012). Fsl. Neuroimage, 62(2), 782–790, doi:10.1016/j.neuroimage.2011.09.015.

Kanwisher, N. (2010). Functional specificity in the human brain: a window into the functional architecture of the mind. Proc Natl Acad Sci U S A, 107(25), 11163–11170, doi:10.1073/pnas.1005062107.

Kiviniemi, V., Starck, T., Remes, J., Long, X., Nikkinen, J., Haapea, M., et al. (2009). Functional segmentation of the brain cortex using high model order group PICA. Hum Brain Mapp, 30(12), 3865–3886, doi:10.1002/hbm.20813.

Laird, A. R., Fox, P. M., Eickhoff, S. B., Turner, J. A., Ray, K. L., McKay, D. R., et al. (2011). Behavioral interpretations of intrinsic connectivity networks. J Cogn Neurosci, 23(12), 4022–4037, doi:10.1162/jocn_a_00077.

Laird, A. R., Lancaster, J. L., & Fox, P. T. (2005). BrainMap: the social evolution of a human brain mapping database. Neuroinformatics, 3(1), 65–78.

Mesulam, M. (2009). Defining neurocognitive networks in the BOLD new world of computed connectivity. Neuron, 62(1), 1–3, doi:10.1016/j.neuron.2009.04.001.

Mesulam, M. M. (1990). Large-scale neurocognitive networks and distributed processing for attention, language, and memory. Ann Neurol, 28(5), 597–613, doi:10.1002/ana.410280502.

Moeller, S., Yacoub, E., Olman, C. A., Auerbach, E., Strupp, J., Harel, N., et al. (2010). Multiband multislice GE-EPI at 7 tesla, with 16-fold acceleration using partial parallel imaging with application to high spatial and temporal whole-brain fMRI. Magn Reson Med, 63(5), 1144–1153, doi:10.1002/mrm.22361.

Pessoa, L. (2014). Understanding brain networks and brain organization. Physics of Life Reviews, 11(3), 400–435, doi:DOI 10.1016/j.plrev.2014.03.005.

Poldrack, R. A. (2006). Can cognitive processes be inferred from neuroimaging data? Trends Cogn Sci, 10(2), 59–63, doi:10.1016/j.tics.2005.12.004.

Poldrack, R. A. (2011). Inferring mental states from neuroimaging data: from reverse inference to large-scale decoding. Neuron, 72(5), 692–697, doi:10.1016/j.neuron.2011.11.001.

Power, J. D., Cohen, A. L., Nelson, S. M., Wig, G. S., Barnes, K. A., Church, J. A., et al. (2011).

Functional network organization of the human brain. Neuron, 72(4), 665–678, doi:10.1016/j.neuron.2011.09.006.

Power, J. D., Schlaggar, B. L., Lessov-Schlaggar, C. N., & Petersen, S. E. (2013). Evidence for Hubs in Human Functional Brain Networks. Neuron, 79(4), 798–813, doi:DOI 10.1016/j.neuron.2013.07.035.

Rubinov, M., & Sporns, O. (2010). Complex network measures of brain connectivity: uses and interpretations. Neuroimage, 52(3), 1059–1069, doi:10.1016/j.neuroimage.2009.10.003.

Sadaghiani, S., & Kleinschmidt, A. (2013). Functional interactions between intrinsic brain activity and behavior. Neuroimage, 80, 379–386, doi:10.1016/j.neuroimage.2013.04.100.

Salimi-Khorshidi, G., Douaud, G., Beckmann, C. F., Glasser, M. F., Griffanti, L., & Smith, S. M. (2014). Automatic denoising of functional MRI data: combining independent component analysis and hierarchical fusion of classifiers. Neuroimage, 90, 449–468, doi:10.1016/j.neuroimage.2013.11.046.

Satterthwaite, T. D., Elliott, M. A., Gerraty, R. T., Ruparel, K., Loughead, J., Calkins, M. E., et al. (2013). An improved framework for confound regression and filtering for control of motion artifact in the preprocessing of resting-state functional connectivity data. Neuroimage, 64, 240–256, doi:10.1016/j.neuroimage.2012.08.052.

Schwarz, G. (1978). Estimating the dimension of a model. The annals of statistics, 6(2), 461–464.

Sepulcre, J., Sabuncu, M. R., Yeo, T. B., Liu, H. S., & Johnson, K. A. (2012). Stepwise Connectivity of the Modal Cortex Reveals the Multimodal Organization of the Human Brain. Journal of Neuroscience, 32(31), 10649–10661, doi:DOI 10.1523/Jneurosci.0759-12.2012.

Setsompop, K., Gagoski, B. A., Polimeni, J. R., Witzel, T., Wedeen, V. J., & Wald, L. L. (2012). Blipped-controlled aliasing in parallel imaging for simultaneous multislice echo planar imaging with reduced g-factor penalty. Magn Reson Med, 67(5), 1210–1224, doi:10.1002/mrm.23097.

Smith, S. M., Beckmann, C. F., Andersson, J., Auerbach, E. J., Bijsterbosch, J., Douaud, G., et al. (2013). Resting-state fMRI in the Human Connectome Project. Neuroimage, 80, 144–168, doi:10.1016/j.neuroimage.2013.05.039.

Smith, S. M., Fox, P. T., Miller, K. L., Glahn, D. C., Fox, P. M., Mackay, C. E., et al. (2009). Correspondence of the brain’s functional architecture during activation and rest. Proceedings of the National Academy of Sciences of the United States of America, 106(31), 13040–13045, doi:DOI 10.1073/pnas.0905267106.

Smith, S. M., Jenkinson, M., Woolrich, M. W., Beckmann, C. F., Behrens, T. E. J., Johansen-Berg, H., et al. (2004). Advances in functional and structural MR image analysis and implementation as FSL. Neuroimage, 23, S208–S219, doi:10.1016/j.neuroimage.2004.07.051.

Sporns, O. (2013). Network attributes for segregation and integration in the human brain. Curr Opin Neurobiol, 23(2), 162–171, doi:10.1016/j.conb.2012.11.015.

Sporns, O. (2014). Contributions and challenges for network models in cognitive neuroscience. Nat Neurosci, 17(5), 652–660, doi:10.1038/nn.3690.

Spreng, R. N., Stevens, W. D., Chamberlain, J. P., Gilmore, A. W., & Schacter, D. L. (2010). Default network activity, coupled with the frontoparietal control network, supports goal-directed cognition. Neuroimage, 53(1), 303–317, doi:10.1016/j.neuroimage.2010.06.016.

Tomasi, D., & Volkow, N. D. (2010). Functional connectivity density mapping. Proc Natl Acad Sci U S A, 107(21), 9885–9890, doi:10.1073/pnas.1001414107.

Tomasi, D., & Volkow, N. D. (2011). Functional connectivity hubs in the human brain. Neuroimage, 57(3), 908–917, doi:10.1016/j.neuroimage.2011.05.024.

van den Heuvel, M. P., & Sporns, O. (2011). Rich-Club Organization of the Human Connectome. Journal of Neuroscience, 31(44), 15775–15786, doi:DOI 10.1523/Jneurosci.3539-11.2011.

Van Essen, D. C., Smith, S. M., Barch, D. M., Behrens, T. E., Yacoub, E., Ugurbil, K., et al. (2013). The WU-Minn Human Connectome Project: an overview. Neuroimage, 80, 62–79, doi:10.1016/j.neuroimage.2013.05.041.

Van Essen, D. C., Ugurbil, K., Auerbach, E., Barch, D., Behrens, T. E., Bucholz, R., et al. (2012). The Human Connectome Project: a data acquisition perspective. Neuroimage, 62(4), 2222–2231, doi:10.1016/j.neuroimage.2012.02.018.

Wig, G. S., Laumann, T. O., & Petersen, S. E. (2014). An approach for parcellating human cortical areas using resting-state correlations. Neuroimage, 93 Pt 2, 276–291, doi:10.1016/j.neuroimage.2013.07.035.

Xu, J., Calhoun, V. D., Worhunsky, P. D., Xiang, H., Li, J., Wall, J. T., et al. (2015). Functional network overlap as revealed by fMRI using sICA and its potential relationships with functional heterogeneity, balanced excitation and inhibition, and sparseness of neuron activity. PLoS One, 10(2), e0117029, doi:10.1371/journal.pone.0117029.

Yarkoni, T., Poldrack, R. A., Nichols, T. E., Van Essen, D. C., & Wager, T. D. (2011). Large-scale automated synthesis of human functional neuroimaging data. Nat Methods, 8(8), 665–670, doi:10.1038/nmeth.1635.

Yeo, B. T., Krienen, F. M., Chee, M. W., & Buckner, R. L. (2013). Estimates of segregation and overlap of functional connectivity networks in the human cerebral cortex. Neuroimage, 88C, 212–227, doi:10.1016/j.neuroimage.2013.10.046.

Yeo, B. T., Krienen, F. M., Eickhoff, S. B., Yaakub, S. N., Fox, P. T., Buckner, R. L., et al. (2014). Functional Specialization and Flexibility in Human Association Cortex. Cerebral Cortex, doi:10.1093/cercor/bhu217.

Yeo, B. T., Krienen, F. M., Sepulcre, J., Sabuncu, M. R., Lashkari, D., Hollinshead, M., et al. (2011). The organization of the human cerebral cortex estimated by intrinsic functional connectivity. J Neurophysiol, 106(3), 1125–1165, doi:10.1152/jn.00338.2011.

Zeki, S. (1993). A vision of the brain. Oxford; Boston: Blackwell Scientific Publications.

Zuo, X. N., Kelly, C., Adelstein, J. S., Klein, D. F., Castellanos, F. X., & Milham, M. P. (2010). Reliable intrinsic connectivity networks: test-retest evaluation using ICA and dual regression approach. Neuroimage, 49(3), 2163–2177, doi:10.1016/j.neuroimage.2009.10.080.

