## Supplementary material for "Characterization of functional diversity of the human brain based on intrinsic connectivity networks"

**Supplemental Fig. 1**

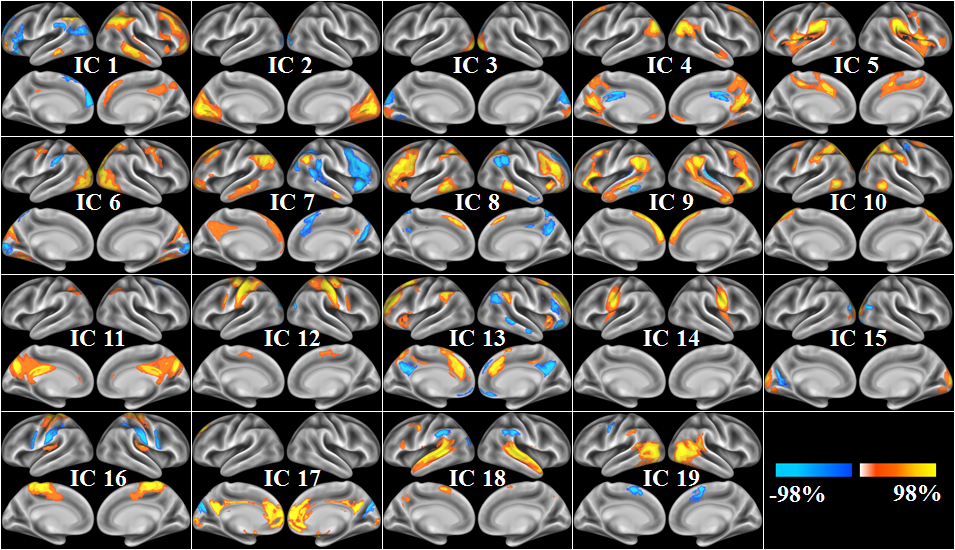

The selected 19 independent components from ICA decomposition with model order of 20. For clearly showing the significant part of each component, the components were transformed with threshold at $\left| Z \right|>5$.

**The selection of model order**

The question about how to decide the optimal model order of ICA remains debated (Beckmann, 2012). There have been some criteria based on information theory to estimate the theoretic optimal model order (Beckmann and Smith, 2004; Calhoun et al., 2002). However, these metrics are difficult to generate the same estimation across subjects and different preprocesses. The biological complexity underlying organization of human brain maybe partly account for the unstable estimation. So far, the various metrics mainly provide the reference, not the decision, about the model order. We still focused on whether the ICA decomposition could provide biologically interpretable components. Smith et.al found that the 20-dimensional decomposition could find the large-scale network structure in resting-state fMRI data (Smith et al., 2009). The high dimensionality (model order of 70) also decomposed biological network structure in a sub-network level. For example, the sensory–motor network was split as separate left and right parts. Kiviniemi et al. considered 70-dimensionality as high model order to segment the brain cortex (Kiviniemi et al., 2009). The default model network was found to be decomposed as the posterior and anterior parts with increment of model order. With a much higher model order of 100, Smith et al. considered each component as a node to detect the organization of whole brain connectome (Smith et al., 2013). Therefore, the high model order could provide a brain segment where the sub-network or even the node of networks emerged. The low dimensionality could detect the large-scale network structures that were similar with the patterns found by seed-based methods. Here, we focused on the large-scale functional properties. We set the model order as 20. From the supplemental Fig. 1, we could observe the well-known resting-state networks, such as default model network (IC4), executive control network (IC1 & IC7) and so on. In fact, we also test our approach across different model order (model order: 30, 40, 50, 60, 70; Supplemental Fig. 2, 3, 4).We found that the results of CoHo index were similar across different model order (Supplemental Fig. 2), and the histograms of CoHo values were still bimodal (Supplemental Fig. 3). We noticed that the primary cortex, such as the sensory-motor cortex, still showed low FD index and the brain regions with high FD index were mainly located in association cortex, such as the precuneus across different model order (Supplemental Fig. 4). In addition, we must point out that the significant brain regions with low or high FD values indeed changed with the increment of model order. It might be caused by the emerging sub-network structures. As the brain was organized hierarchically, different properties might be correlated with the particular hierarchical level.

**Supplemental Fig. 2**

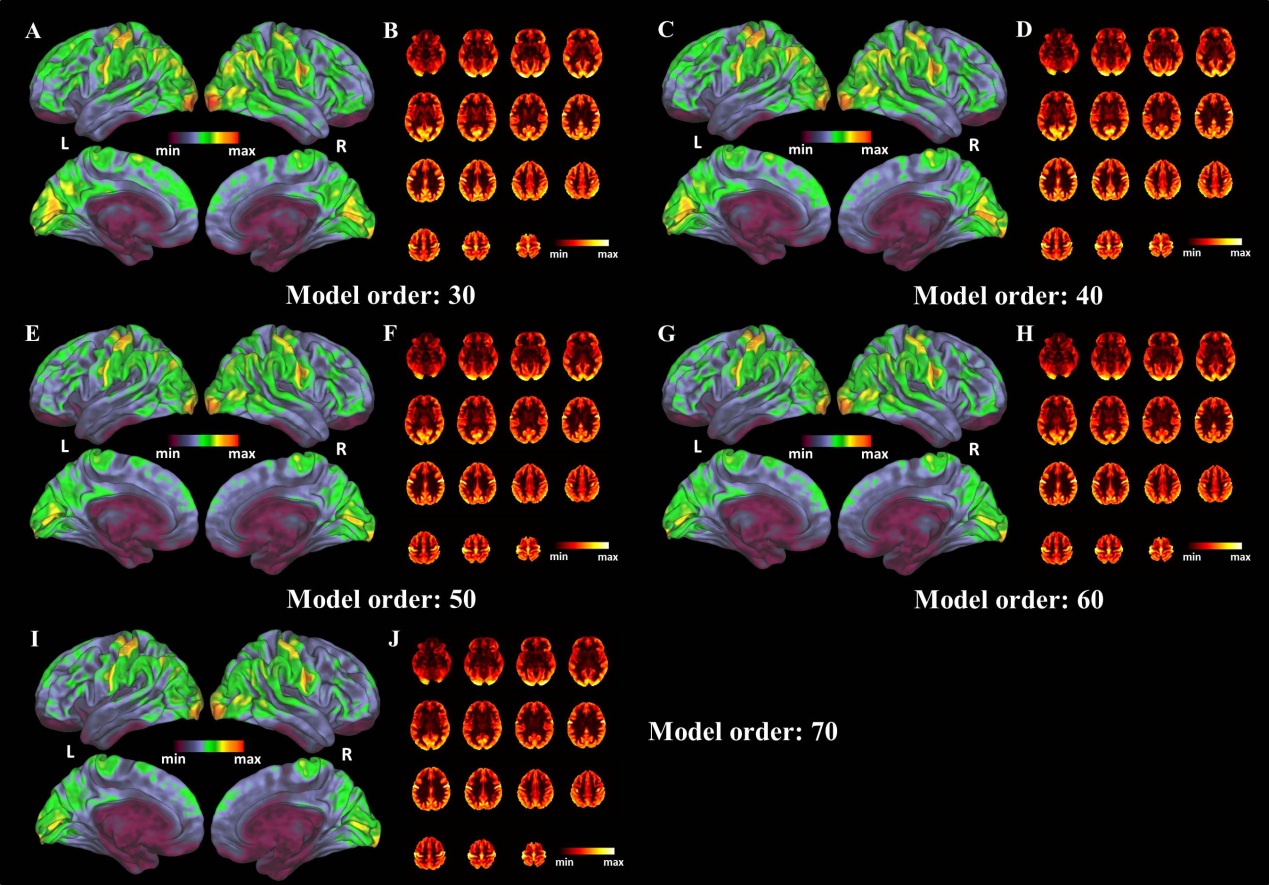

The results of CoHo index across different model order.

**Supplemental Fig. 3**

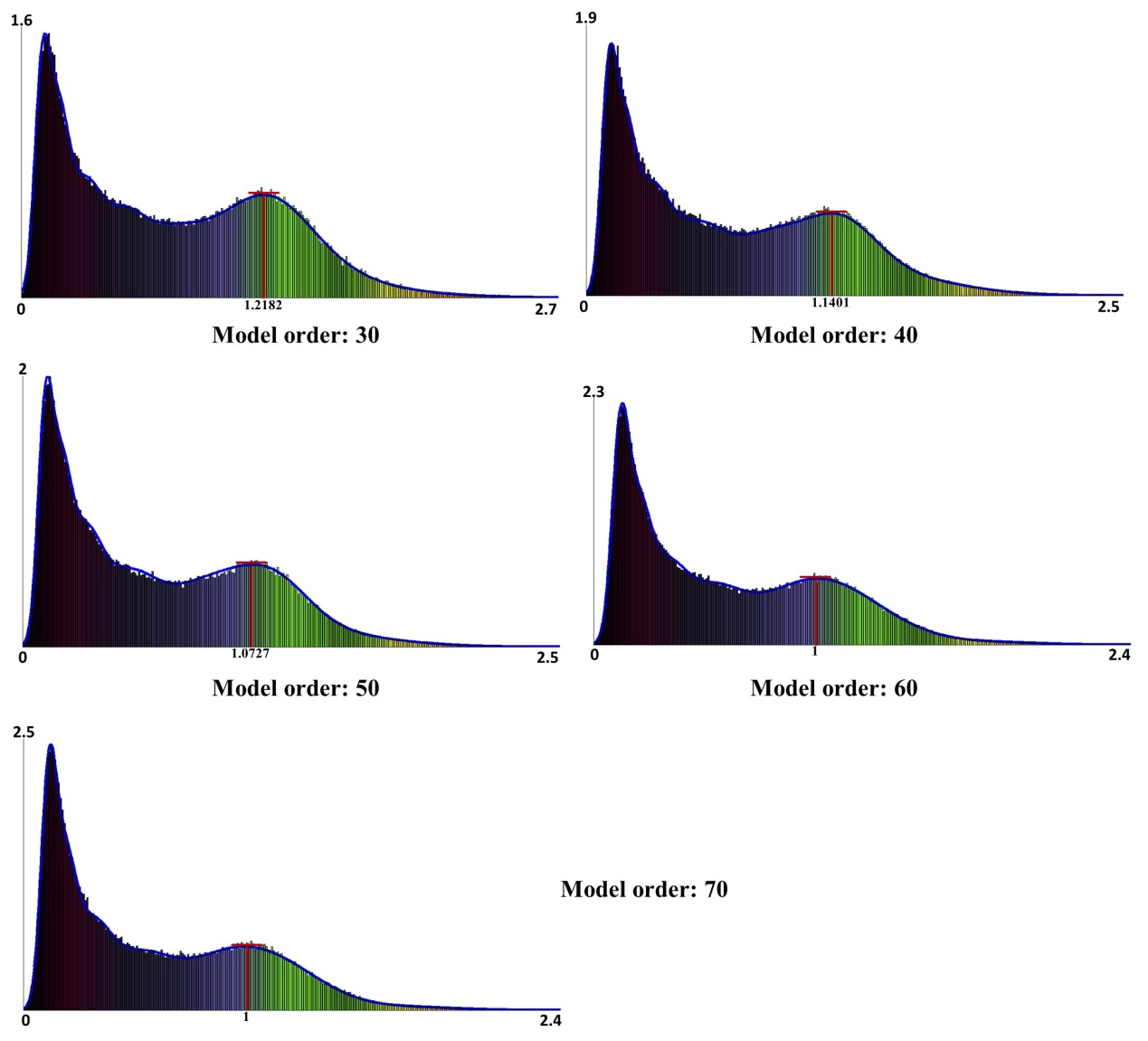

The histograms of CoHo index across different model order. The red lines were corresponding with the second peak point in the estimated density functions, which were used as the threshold to deduce the mask for the next-step calculation of FD index.

**Supplemental Fig. 4**

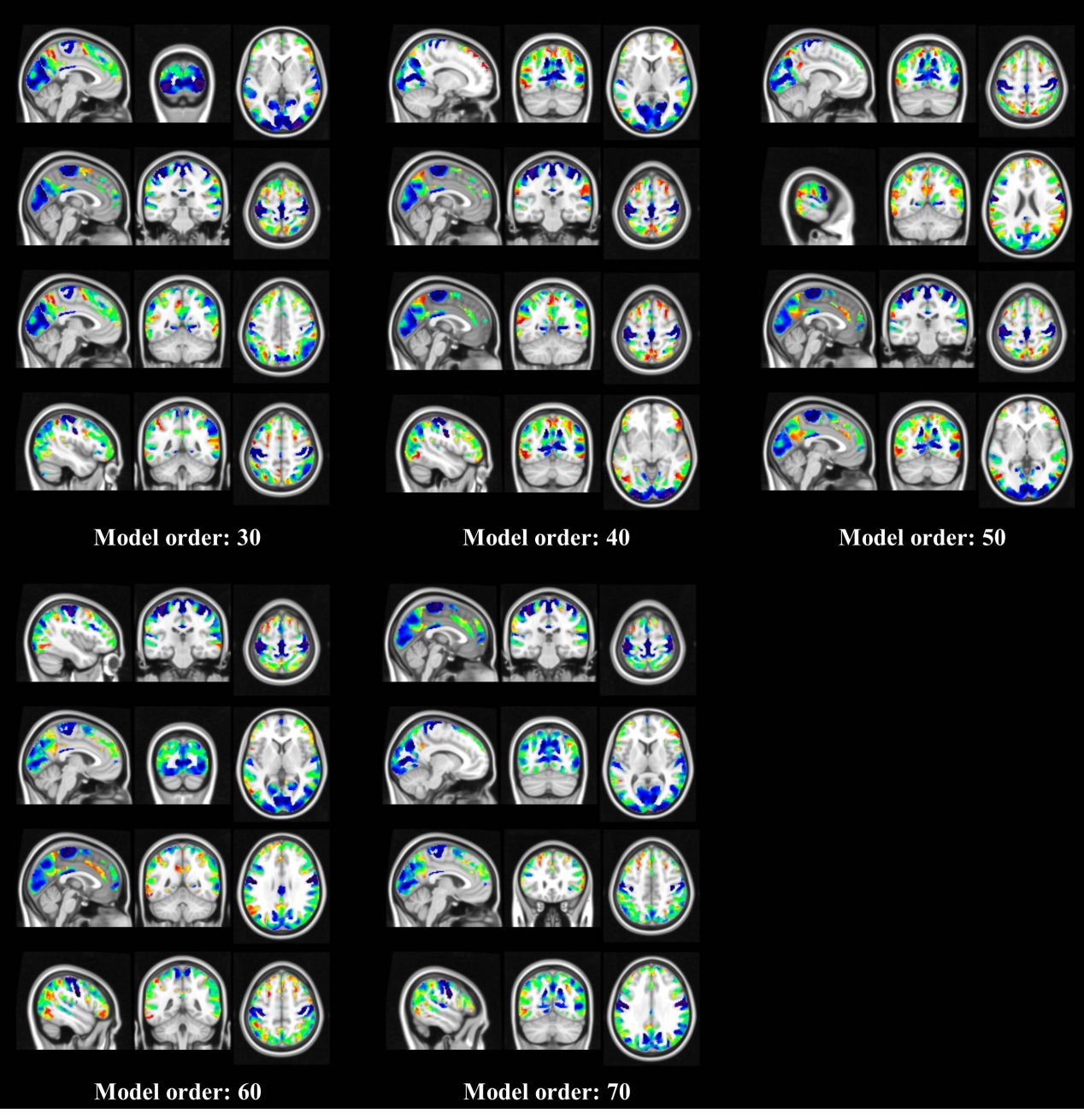

The results of FD index calculated in the CoHo-deduced masks.

**The calculation of subject-level FD index**

The subject-level FD index was proposed to be calculated based on the ICA with reference. In detail, the approach named Multivariate Objective Optimization ICA with Reference (MOO-ICAR) was adopted (Du and Fan, 2013). The components derive from the group were used as the reference to calculate the subject-level ICA-decomposition. Two optimizations were considered by a multi-objective optimization strategy that optimized the similarity between the group components and the deduced subject-level components and the independence between subject-level components. After the subject-level ICA-decomposition was deduced, we calculated the CoHo index and FD index on each subject. We gave an overview of the analysis (Supplemental Fig. 5).

**Supplemental Fig. 5**

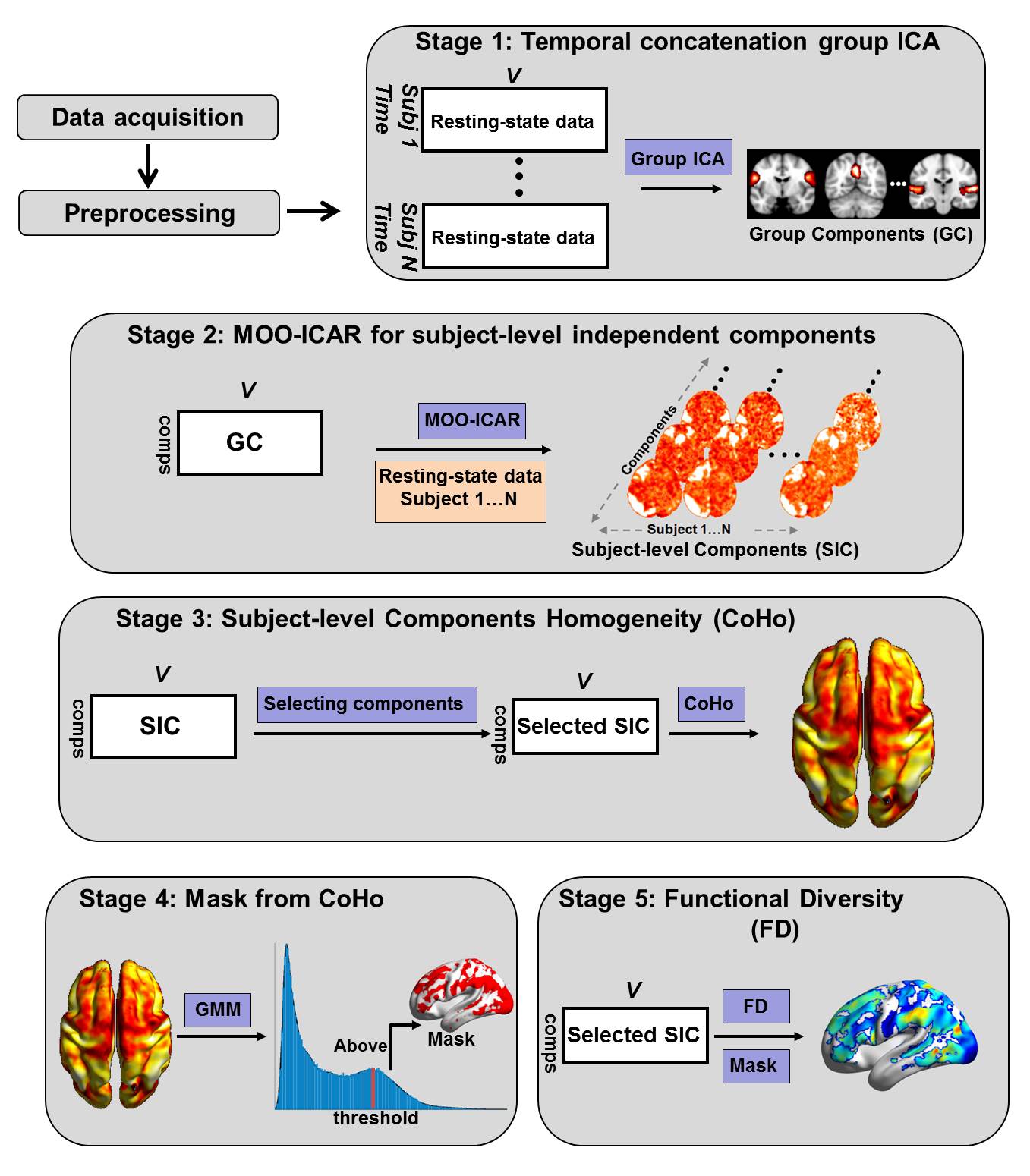

An overview about the analysis of subject-level FD.

We showed the results of CoHo and FD index on a randomly selected subject (CoHo result: Supplemental Fig. 6; FD result: Supplemental Fig. 7). We noticed that the histogram of subject-level CoHo index did not show the bimodal distribution (Supplemental Fig. 6A). It might be caused by the low signal-noise ratio in subject level. However, we could still observe the trend that the latter part of the histogram was slowly changed. Using the GMM model, we modeled the probability density function of CoHo index. We used the second derivative of the modeled probability density function to find the point corresponding with the slowest change in the latter part (Supplemental Fig. 6B). We just gave an approach to deduce the subject-level threshold for the CoHo index. Other approach could also be tried. Further, we found that the distribution of subject-level FD index, especially the significant brain regions with high / low FD values (at top / bottom 5% and cluster-size > 80 voxels), was similar with the group-level result (Supplemental Fig. 7).

**Supplemental Fig. 6**

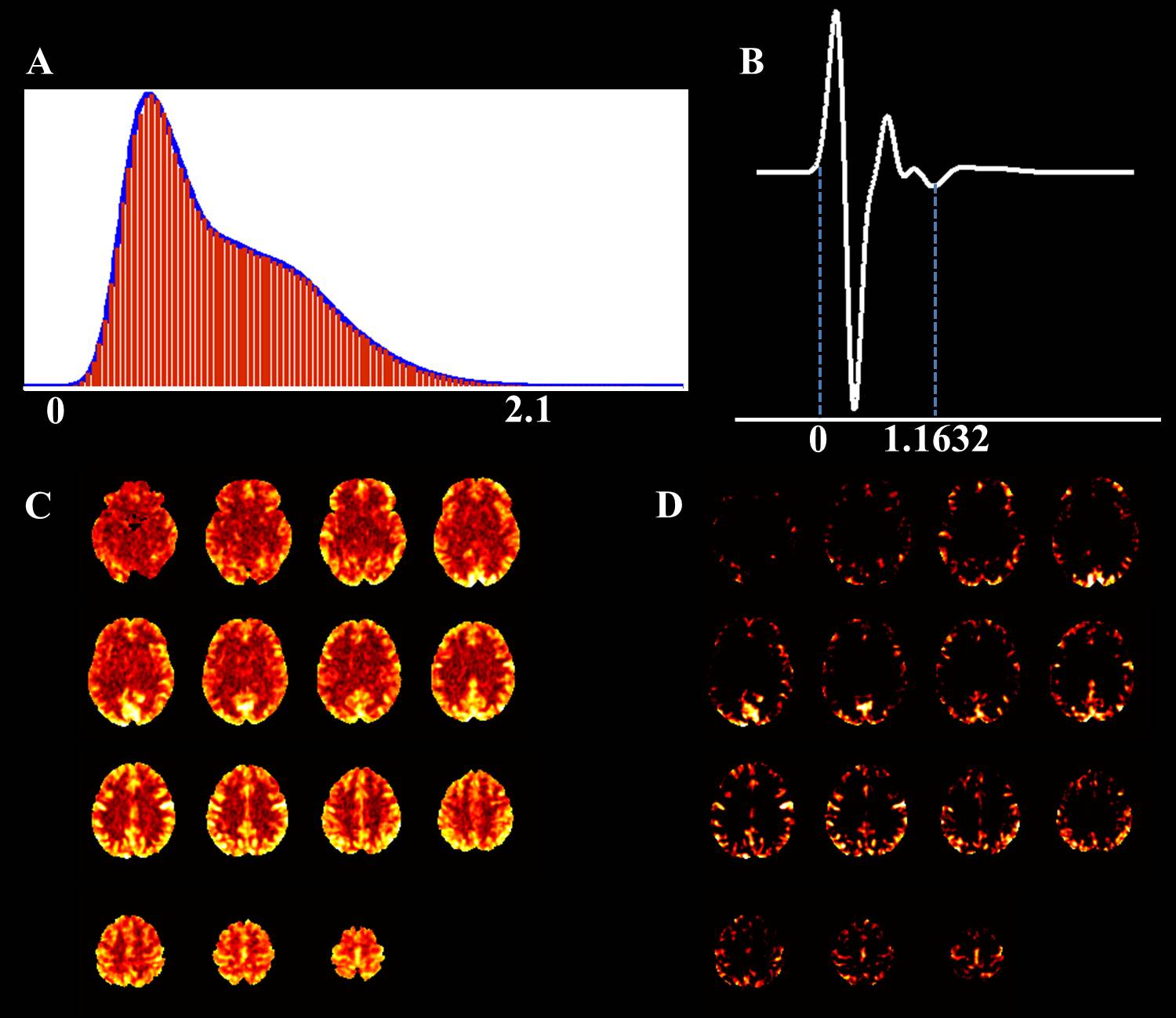

The subject-level CoHo map and the GMM estimation. (A) The estimation of GMM. The blue curve was the Gaussian mixture function to model the underlying probability density function. The red histogram was derived from the CoHo map. (B) The second derivative of the modeled probability density function. The latter blue dash line was the chosen threshold, which represented the slowest changed point in the latter part of the function. (C) The transverse slices of subject-level CoHo map. (D) The transverse slices of the brain regions with CoHo values greater than the threshold.

**Supplemental Fig. 7**

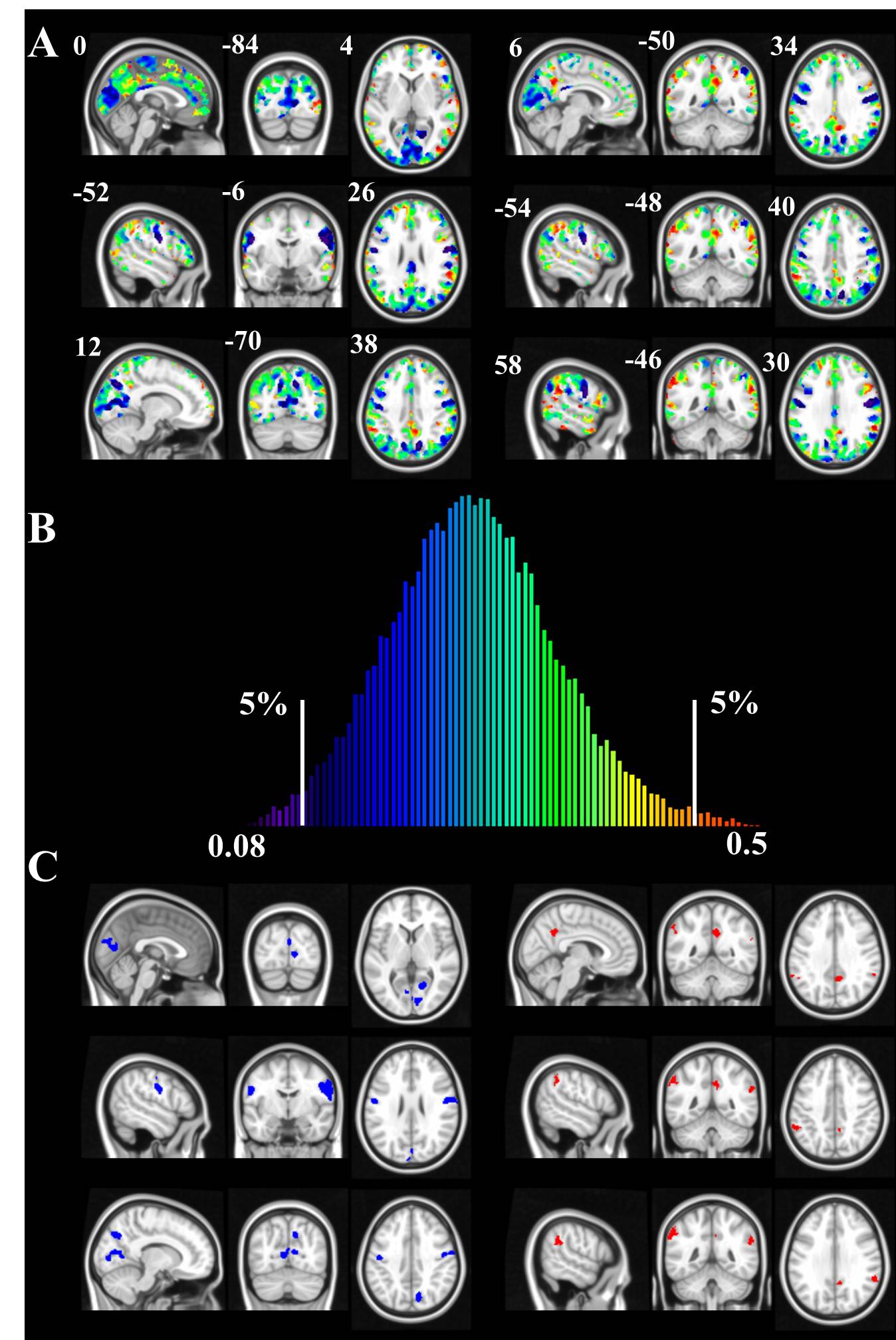

The distribution of FD index. (A) The derived FD index within mask. (B) The histogram of FD index. The color in the histogram was corresponding to the color on (A). (C) The top / bottom 5% of brain regions with high / low FD values. The cluster size was larger than 80 voxels. The blue regions in left side were corresponding to low FD values. The red regions in right side were corresponding to high FD values.

**Reference:**

Beckmann, C.F., 2012. Modelling with independent components. Neuroimage 62, 891-901.

Beckmann, C.F., Smith, S.A., 2004. Probabilistic independent component analysis for functional magnetic resonance imaging. Ieee Transactions on Medical Imaging 23, 137-152.

Calhoun, V.D., Adali, T., Pearlson, G.D., Pekar, J.J., 2002. A method for making group inferences from functional MRI data using independent component analysis. Hum Brain Mapp 16, 131-131.

Du, Y., Fan, Y., 2013. Group information guided ICA for fMRI data analysis. Neuroimage 69, 157-197.

Kiviniemi, V., Starck, T., Remes, J., Long, X., Nikkinen, J., Haapea, M., Veijola, J., Moilanen, I., Isohanni, M., Zang, Y.F., Tervonen, O., 2009. Functional segmentation of the brain cortex using high model order group PICA. Hum Brain Mapp 30, 3865-3886.

Smith, S.M., Fox, P.T., Miller, K.L., Glahn, D.C., Fox, P.M., Mackay, C.E., Filippini, N., Watkins, K.E., Toro, R., Laird, A.R., Beckmann, C.F., 2009. Correspondence of the brain's functional architecture during activation and rest. Proceedings of the National Academy of Sciences of the United States of America 106, 13040-13045.

Smith, S.M., Vidaurre, D., Beckmann, C.F., Glasser, M.F., Jenkinson, M., Miller, K.L., Nichols, T.E., Robinson, E.C., Salimi-Khorshidi, G., Woolrich, M.W., Barch, D.M., Ugurbil, K., Van Essen, D.C., 2013. Functional connectomics from resting-state fMRI. Trends Cogn Sci 17, 666-682.
